# Poldip2 deficiency attenuates disease severity in a mouse model of COVID-19

**DOI:** 10.1101/2025.06.17.657579

**Authors:** Ruinan Hu, Alejandra Valdivia, Taylor White, Willy Ju, Maegan L. Brockman, Zhan Zhang, Hongyan Qu, Georgette Gafford, Giji Joseph, Samantha Burton, Leda Bassit, Tysheena P. Charles, Rebecca D. Levit, Cynthia A. Derdeyn, Kathy K. Griendling, Bernard Lassègue, Marina S. Hernandes

**Author notes:** Corresponding author Marina S. HernandesEmory University School of Medicine Department of Medicine, Division of Cardiology 1750 Haygood Dr. NE Atlanta, GA 30322, USA Phone: [+1].

## Abstract

The lungs are the primary target of severe acute respiratory syndrome coronavirus 2 (SARS-CoV- 2), with the infection resulting in lung inflammation, pulmonary vascular leakage and diffuse alveolar damage. Polymerase delta-interacting protein-2 (Poldip2) mediates lung inflammation and vascular permeability after lipopolysaccharide-induced acute respiratory distress syndrome; however, its role in regulating lung permeability, vascular inflammation and tissue damage following SARS-CoV-2 infection is completely unknown. Here, we assessed the role of Poldip2 in inflammation, immune cell infiltration and lung tissue damage in response to SARS-CoV-2 infection. Our data shows that while deletion of Poldip2 does not affect the susceptibility to SARS-CoV-2 infection, mice heterozygous for Poldip2 exhibit reduced lung tissue damage, reduced cytokine and chemokine induction and decreased infiltration of myeloperoxidase (MPO)-positive neutrophils into inflamed lung tissue. These data reveal that Poldip2 depletion mitigates inflammation and immune cell infiltration following SARS-CoV-2 infection, highlighting the therapeutic potential of Poldip2 inhibition to attenuate severe lung injury.

## Introduction

Severe acute respiratory syndrome coronavirus-2 (SARS-CoV-2) initiated the global Coronavirus Disease 2019 (COVID-19) pandemic. According to the World Health Organization, as of January 2025, more than 750 million SARS-CoV-2 identified cases have been reported with over 7 million confirmed deaths [1]. SARS-CoV-2 is primarily transmitted through respiratory droplets and infects human cells via binding to angiotensin-converting enzyme 2 (ACE2), which is the main SARS-CoV-2 receptor [2]. Early in infection, SARS-CoV-2 has been shown to target nasal and bronchial epithelial cells and pneumocytes, resulting in high copy numbers in the lower respiratory tract [3]. Pro-inflammatory signaling molecules are released by infected cells and alveolar macrophages, which then trigger the recruitment of T lymphocytes, monocytes, and neutrophils, contributing to further aggravating the inflammatory response. In the late stages of infection, damage to vascular endothelial cells and alveolar epithelial cells compromises the epithelial-endothelial barrier integrity, leading to vascular leakage and pulmonary edema [3], with accumulation of fluid containing large amounts of proteins, cellular debris, hyaline membrane formation [4–7] and early-phase acute respiratory distress syndrome [3].

Additional pathophysiological mechanisms associated with SARS-CoV-2 infection include an acute inflammatory response which was found to be positively correlated with the severity of COVID symptoms [8, 9]. Evaluation of bronchoalveolar lavage (BAL) fluid collected from COVID- 19 patients revealed high levels of pro-inflammatory cytokines including IL-1β, IL-8 and TNFα [10], suggesting that lung inflammation is a major component of SARS-CoV-2 pathogenesis. Corroborating these findings from human samples, SARS-CoV-2 infection has been shown to induce TNFα, IFN-γ, MCP1 and IL-6 protein expression in the lung tissue of humanized ACE2 transgenic mice [11]. In a different study, a liquid chromatography-mass spectrometry-based proteomic and phosphoproteomic evaluation of lung tissue samples isolated from SARS-CoV-2 infected K18-hACE2 mice showed the presence of hyperphosphorylated proteins associated with leukocyte transendothelial migration [12]. The same study also demonstrated enhanced phosphorylation of tight junction proteins in lung tissue in response to SARS-CoV-2 infection [12], which is a critical mechanism involved in endothelial barrier dysfunction [13] and acute lung injury [4].

Polymerase δ-interacting protein 2 (Poldip2) was originally identified as a binding partner of polymerase-δ and was proposed to be involved in DNA damage repair [14, 15]. In more recent studies, our lab discovered that Poldip2 is a critical regulator of inflammation and endothelial barrier function [13, 16–18]. Using a lipopolysaccharide (LPS)-induced acute respiratory distress syndrome (ARDS) model, our group has recently reported that Poldip2 knockdown decreases cytokine and chemokine induction and lung immune cell infiltration [18, 19]. Our *in vitro* studies further demonstrated that Poldip2 downregulation in pulmonary microvascular endothelial cells attenuates the protein expression of leukocyte adhesion molecules and decreases LPS-induced THP-1 monocyte adherence [19], suggesting that Poldip2 plays a role in lung endothelial cells during inflammation. In subsequent studies we demonstrated that Poldip2 depletion in pulmonary microvascular endothelial cells prevents disruption of VE-cadherin, one of the main endothelial cell junction proteins, and the subsequent increase in permeability in response to TNFα, likely in a Rho pathway-dependent manner [18]. Taken together, these results suggested an important role for Poldip2 in inflammation and endothelial barrier function during ARDS.

Based on these observations, we hypothesized that depletion of Poldip2 would also be protective against SARS-CoV-2-induced lung injury. To test this hypothesis, we crossed K18- humanized ACE2 transgenic mice with Poldip2^+/-^ mice. Following intranasal inoculation with SARS-CoV-2, disease severity, lung tissue damage, inflammation and immune cell infiltration were evaluated. While heterozygous deletion of Poldip2 does not affect the susceptibility to SARS-CoV-2 infection, our data indicate that Poldip2^+/-^ mice exhibit reduced lung tissue damage, cytokine and chemokine induction and decreased infiltration of myeloperoxidase (MPO)-positive neutrophils. These data reveal that Poldip2 depletion mitigates inflammation and immune cell infiltration following SARS-CoV-2 infection, highlighting the therapeutic potential of Poldip2 inhibition indicating that targeted inhibition of Poldip2 may be an option for therapeutic intervention moving forward.

## Materials and Methods

### Animals

K18-hACE2 transgenic (strain # 034860) and C57BL/6J (strain # 000664) mice were purchased from The Jackson Laboratory. Genetic drift was minimized by backcrossing K18- hACE2 to C57BL/6. Genotyping was performed either using a traditional 4 primer PCR assay (**Table 1**), or by real-time PCR when it was necessary to detect homozygotes. In the latter case, the assay took advantage of an SNP adjacent to the ACE2 transgene. It was performed with a QuantStudio 7 instrument (Thermo Fisher), Luna Universal Probe qPCR Master Mix (New England Biolabs) and two primers spanning 220 bp. Wild type and mutant alleles were respectively detected using FAM and HEX labeled TaqMan probes (Millipore Sigma) (**Table 1**). Annealing and extension were carried out at 58°C for 40 seconds. Constitutive Poldip2 knockout mice were produced and genotyped as described previously [20] or using a real-time PCR assay based on high-resolution melting curve analysis (**Table 1**). Experimental animals were generated by crossing homozygous ACE2 transgenics with Poldip2^+/-^ mice to obtain equal numbers of heterozygous ACE2 transgenics with either Poldip2^+/-^ or Poldip2^+/+^ (littermate controls) genotypes. In all experiments, male and female human ACE2 and Poldip2 heterozygous mice and littermate controls aged 3-5 months were randomly assigned to control and experimental groups using a random number generator. The number of female and male animals in each group was approximately similar. This study was carried out in compliance with the ARRIVE guidelines [21] and all animal welfare and experimental protocols were approved by the Emory University Institutional Animal Care and Use Committee. Ethical approval was received before conducting the study. Researchers performing analyses were blinded to genotype and experimental groupings of mice.

**Table 1:**
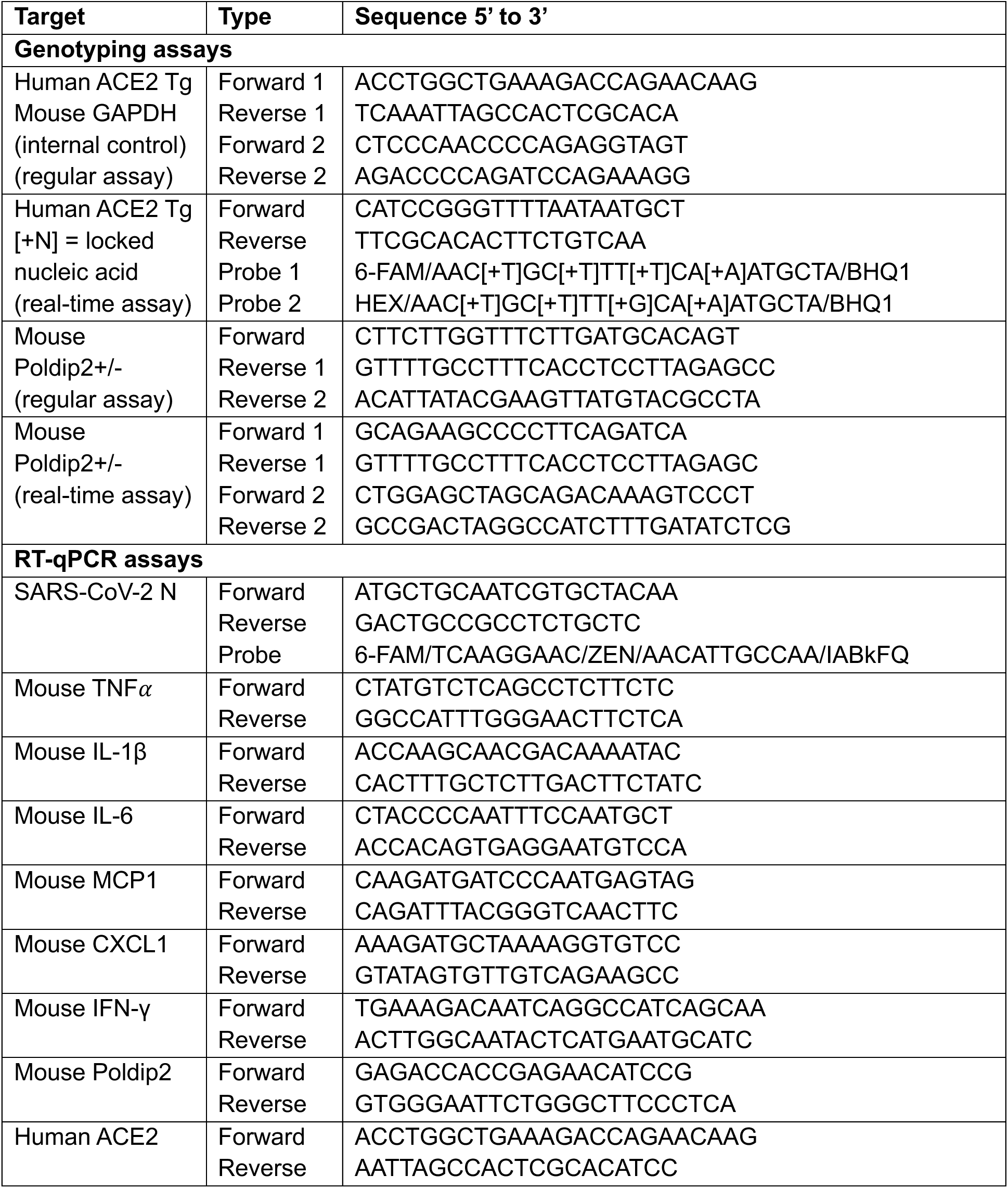
Primer and probe sequences

### Virus preparation

The SARS-CoV-2 strain (hCoV-19/US-WA1/2020) was obtained from Biodefense and Emerging Infections Research Resources Repository (Cat NR53899, Lot 7004383, BEI Resources). The virus was propagated in Vero E6/TMPRSS2 cells (passage 5, from Japanese Collection of Research Bioresources Cell Bank, Sekisui XenoTech, LLC) under 37°C/5% CO_2_, the multiplicity of infection (MOI) was 0.025. The virus was harvested after 48 hours, or when 80% cytopathic effect was observed. Virus titer was determined by 50% tissue culture infective dose (TCID_50_/ml) or plaque assays (plaque for PFU/ml). The concentration of the virus was 1.78x10^9^ TCID_50_ (calculated using the Spearman Karber method) and 1.25x10^9^/ml. Small aliquots of virus were made and stored at -80°C or liquid nitrogen until use. All experiments involving infectious virus were conducted at Emory University in approved biosafety level 3 (BSL) laboratories with routine medical monitoring of laboratory personnel.

### Animal model of SARS-CoV-2 infection

Female and male K18-hACE2 transgenic C57BL/6J mice were anesthetized with 3% isoflurane and intranasally infected with 4x10^5^ plaque-forming units (PFU) SARS-CoV-2 virus diluted in phosphate-buffered saline (PBS) in an Animal Biosafety Level 3 (ABSL3) facility at the Emory National Primate Research Center. Uninfected control mice were inoculated with equal volume of PBS. Mice were maintained in Sealsafe HEPA-filtered air in/out units and monitored daily for morbidity, and clinical signs of disease. Body temperature was measured using a non-contact infrared thermometer under light anesthesia (1–1.5% isoflurane) and body weight was recorded every other day. Lung tissue and bronchoalveolar lavage (BAL) fluids were collected 7 days postinfection for determination of inflammatory gene expression, inflammatory protein expression, leucocyte infiltration and histological analysis.

### Determination of *in vivo* viral load in lung tissue

Lung viral load was determined at day 7 postinfection. Briefly, the lower lobes of the mouse lung were homogenized in 1 mL of Trizol reagent (Cat 79306, Qiagen) using one stainless steel bead (3/16" Inch 440 Stainless Steel Ball Bearings G100, BC Precision), in a TissueLyser LT (Cat 85600, Qiagen) at 25 Hz for 10 min. Total RNA was purified with the RNeasy Plus kit (Qiagen). Expression of the SARS-CoV-2 N gene was measured by RT-qPCR using two primers and a TaqMan probe described by Hassan et al [22] (**Table 1**) and purchased from IDT. The assay was carried out with a QuantStudio 7 instrument (Thermo Fisher), using Luna Universal Probe qPCR Master Mix (New England Biolabs) with annealing and extension at 60°C. A cDNA standard of viral gene N was purchased from IDT. Data quantification was performed using the qpcR software library in the R environment, as described in the RT-qPCR section.

### Bronchoalveolar lavage collection

After euthanasia, mouse tracheas were exposed through a small skin incision on the anterior neck and cannulated using a 21-gauge lavage needle. For assessment of cytokine and chemokines, 1.2 ml of PBS was instilled in the tracheal lavage needle and retrieved. Return volume was centrifuged (300 g, 10 min), and the supernatant was incubated in a pre-warmed dry heat block at 56°C for 30 min to inactivate the virus and subsequentially stored at −80°C for assessment of cytokine and chemokines levels by ELISA.

### Immunohistochemistry and immunofluorescence

Human lung samples from the Georgia Medical Examiner’s Office were fixed in 10% formalin in saline for one week, dehydrated using a HistoCore PEARL automatic tissue processor (Leica), transferred to 70% ethanol, embedded in paraffin and sectioned at 5 µm thickness. Following deparaffinization and antigen retrieval using citrate buffer (pH 6.0) in a pressure cooker for 10 min, sections were blocked with 2% IgG-free BSA and 5% normal goat serum in PBS for 1 hour at room temperature. Primary antibodies against Poldip2 (rabbit, HPA007700, Sigma Aldrich) and CD31 (mouse, Ab9498, Abcam), or with the corresponding IgG isotype controls (ab18450 and ab172730, Abcam) were incubated overnight at 4°C. After washing with PBS, sections were incubated with anti-rabbit Alexa FluorTM 568 (Invitrogen A-11011) and anti-mouse Alexa FluorTM 488 (Invitrogen, A-11001) secondary antibodies for 1 hour at room temperature. Nuclei were counterstained with DAPI, and slides were mounted using Diamond Antifade (P36961, Thermo Fisher Scientific).

Images were acquired as z-stack with a Zeiss LSM800 Airyscan laser scanning confocal microscope with a Plan-Apo 63x/1.4 NA objective. Staining, imaging, and quantification were performed by two blinded investigators using ImageJ software. In brief, maximum intensity projections were generated from the z-stacks and used to create a mask with the CD31 signal (green channel) then the mean intensity of Poldip2 staining (red signal) contained in the CD31 area was calculated. Results show the integrated density of Poldip2 staining (mean gray values x pixel number ± standard error of the mean [SEM]).

For mouse tissue processing, after euthanasia, lungs were instilled with 10% formalin through the trachea, excised and placed in 10% formalin for one week. Lungs were then transferred to 70% ethanol, dehydrated using a HistoCore PEARL automatic tissue processor (Leica), paraffin-embedded, and sectioned. Lung sections, 5-μm in thickness, were either stained with hematoxylin (ab220365, Abcam) and eosin (ab246824, Abcam) for morphological analysis or processed for immunofluorescence.

Neutrophil infiltration in mouse lungs was assessed by immunofluorescence following co-staining for the leukocyte-common antigen, CD45, and myeloperoxidase (MPO). Lung sections were covered with water and treated with UV light for 30 min in a humidity chamber to reduce autofluorescence. Antigen retrieval was performed using 10 mM citrate buffer (pH 6.0) in a pressure cooker for 10 minutes. Blocking was carried out for 1 hour using 10% goat serum and 2% IgG-free BSA in Universal Buffer (0.05 M Tris HCl, pH 7.6, 0.15 M NaCl, and 0.1 % sodium azide). Consecutive sections were then incubated overnight at 4°C with primary antibodies: rabbit anti-CD45 (ab10558, Abcam) and rat anti-MPO (ab300650, Abcam), or with the corresponding IgG isotype controls (rabbit ab172730, Abcam and rat ab18450, Abcam), dissolved in blocking buffer. Next, the sections were incubated for 1 hour at room temperature with secondary antibodies: anti-rabbit Alexa Fluor 568 (A11011, Invitrogen), anti-rat Alexa Fluor 488 (A11006, Invitrogen), and DAPI (D9542, Sigma-Aldrich). To further reduce autofluorescence, samples were then incubated with 0.1% Sudan Black in 70% Ethanol for 10 minutes, washed three times with Universal Buffer, and mounted using ProLong Diamond Antifade (P36961, Thermo Fisher Scientific). At least 12 images from each slide were acquired as z-stacks using a Zeiss LSM800 Airyscan laser scanning confocal microscope with an EC Plan-Neofluar 40x/1.3 NA objective. Staining, imaging, and quantification were performed by two blinded investigators. In brief, maximum intensity projections were generated from the z-stacks and used to quantify the number of positive cells with the "Cell Counter" plugin in ImageJ software, as well as to measure the tissue area analyzed in each image. The numbers of MPO and CD45 double positive cells were normalized to total tissue area.

### RNA extraction and RT-qPCR

Mouse lungs were homogenized in 1 mL of QIAzol reagent (Cat 79306, Qiagen) using one stainless steel bead, in a TissueLyser LT (Cat 85600, Qiagen) at 25 Hz for 10 min. Total RNA was purified with the RNeasy Plus kit (Qiagen) or Direct-zol DNA/RNA Miniprep (Cat R2080 Zymo Research). Reverse transcription was performed using Protoscript II reverse transcriptase (Cat M0368, New England Biolabs) with random primers. cDNA was amplified with 2X Forget-Me-Not EvaGreen qPCR Master Mix with Low ROX (Cat 31045, Biotium) and primers against mouse TNF𝛼, IL-1β, IL-6, MCP1, CXCL1, IFN-γ, Poldip2 and human ACE2 (**Table 1**). Reactions were carried out in 96 wells qPCR plates, using a QuantStudio 7 Flex System (Thermo Fisher Scientific) Real-Time qPCR System. Data analysis was performed using the mak3i module of the qpcR software library [23, 24] in the R environment [25]. Data were additionally processed using the NORMA-Gene software [26].

### ELISA

TNF𝛼, IL-1B, IL-6, MCP1, CXCL1, IFN-γ levels in BAL fluids were measured using specific ELISA kits according to the manufacturer’s instructions (R&D Systems).

## Data analysis and statistics

Data are presented as mean ± SEM. A Log transformation was applied when required to normalize distributions, as shown by the Log scale of the Y axis in bar graphs. Human data were analyzed using a two-tailed nested t-test. Mouse data were analyzed using a two-tailed unpaired t*-*test for single comparisons or two-way analysis of variance (ANOVA), followed by the Sidak test for multiple comparisons. A threshold of P< 0.05 was considered significant. Calculations were performed using Prism 10 (GraphPad).

## Results

### Poldip2 is upregulated in human lung after SARS-CoV-2 infection

To begin exploring a possible contribution of Poldip2 to COVID-19 pathology, we analyzed lung tissue sections from control and virus-infected patients. Since our previous results pointed to a positive effect of Poldip2 on vascular endothelial permeability [13, 16, 18, 19, 27], we assessed Poldip2 co-expression with the endothelial marker CD31 by immunofluorescence. As shown in **Figure 1**, endothelial Poldip2 was upregulated in patients with SARS-CoV-2 infection, compared to controls. This observation suggests that one of the ways in which the virus may harm patients is via upregulation of Poldip2, which may in turn aggravate disease by increasing endothelial permeability in lung. Therefore, in the present study we investigated the potential benefit of knocking down Poldip2 in an animal model of COVID-19.

**Figure 1:**
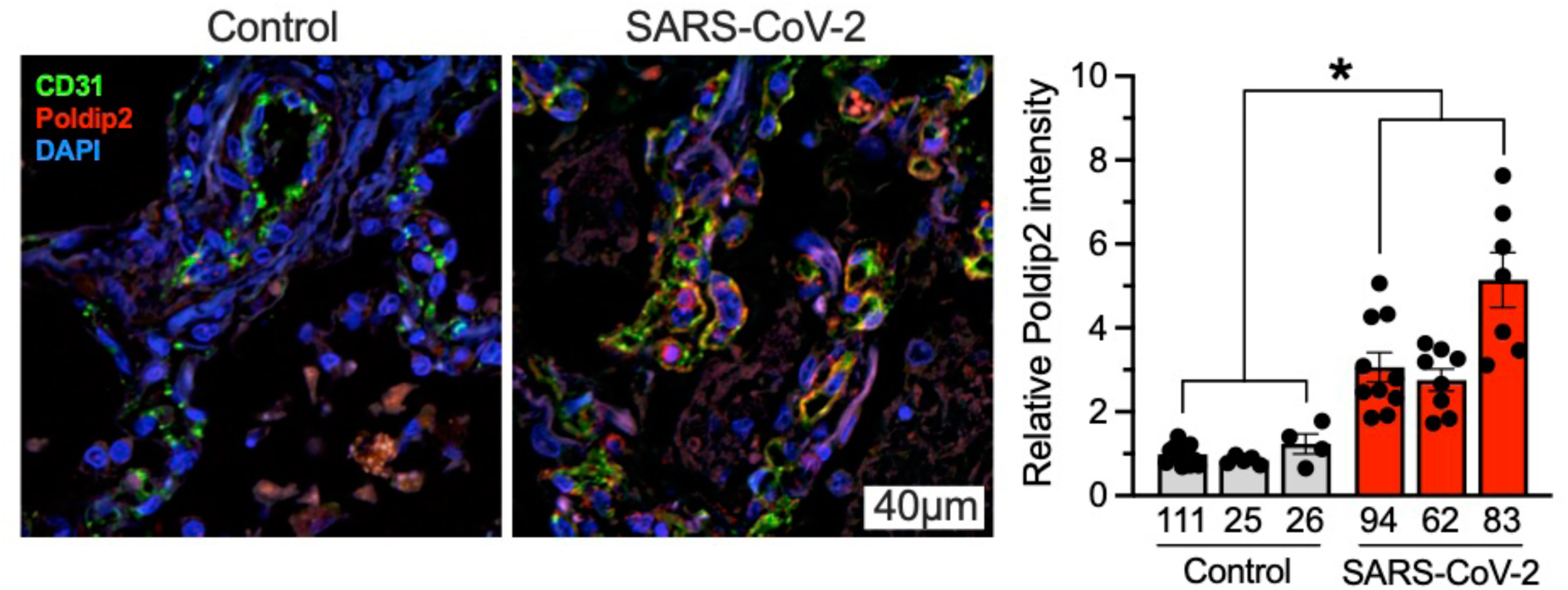
Endothelial Poldip2 is upregulated in human lung by SARS-CoV-2 infection. Sections of lung tissue were immunostained for CD31 (green), Poldip2 (red) and nuclei (blue). Left: representative images from one control and one infected patient. Right: quantification of Poldip2 staining in CD31-positive areas. Bars represent means ± SEM from 4-10 pictures per sample taken at random locations. Data were analyzed using a two-tailed nested Student t-test, n=3, *P<0.05.

### Heterozygous deletion of Poldip2 does not affect the susceptibility to SARS-CoV-2 infection

The first question was whether human ACE2 transgenic, Poldip2^+/+^ or Poldip2^+/-^ (hACE2/Poldip2) mice are equally susceptible to SARS-CoV-2 infection. Mice were infected intranasally with 4x10^5^ PFU SARS-CoV-2 virus. Viral copies in the lungs were measured by RT-PCR on day 7 post infection. As shown in **Figure 2**, no significant difference was found in lung viral RNA expression between Poldip2^+/+^ and Poldip2^+/-^ mice, indicating that Poldip2 knockdown does not affect viral load and, therefore different degrees of susceptibility to SARS-CoV-2 infection would not explain any observed differences in response to the virus. We also confirmed that Poldip2 mRNA levels were decreased by about 50% in Poldip2^+/-^, compared to Poldip2^+/+^ mice (**Supplemental Figure 1A**) and that the expression of human ACE2 mRNA was not affected by Poldip2 depletion (**Supplemental Figure 1B**).

**Figure 2:**
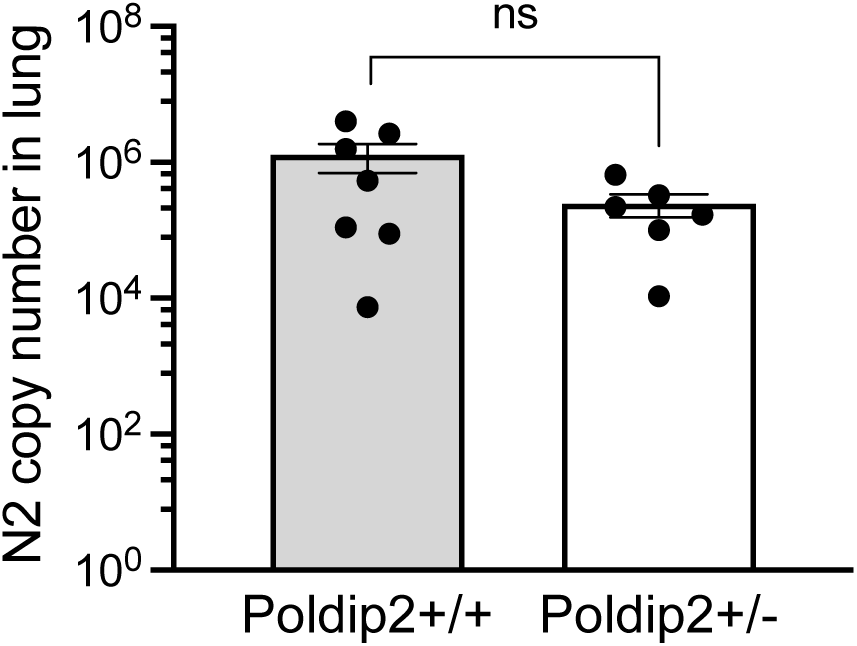
Viral loads are not affected by the Poldip2 genotype. Male or female human ACE2 transgenic, Poldip2^+/+^ or Poldip2^+/-^ (hACE2/Poldip2) mice were infected with SARS-CoV-2 by nasal inoculation. Lungs were collected 7 days later for RNA preparation. Viral loads were measured by RT-qPCR using a primer pair and a TaqMan probe specific for the N2 viral gene. Copy numbers were calculated from a standard curve of viral cDNA. Bars represent means ± SEM of data from n=6-7 animals. Means were compared using an unpaired two-tailed Student t test: ns, not significant.

To further characterize the susceptibility of Poldip2^+/-^ animals to SARS-CoV-2 infection, mice infected intranasally with SARS-CoV-2 were monitored every other day for body weight and temperature changes until day 6 post-infection. As indicated in **Figure 3**, SARS-CoV-2 tended to decrease body temperature in both Poldip2^+/+^ and Poldip2^+/-^ mice groups at day 6 post-infection; however, no significant difference was observed between the two genotypes. Similarly, SARS-CoV-2 infection did not induce a significant weight loss in either Poldip2^+/+^ or Poldip2^+/-^ mice. These data suggest that Poldip2^+/+^ and Poldip2^+/-^ mice are equally susceptible to SARS-CoV-2 infection.

**Figure 3:**
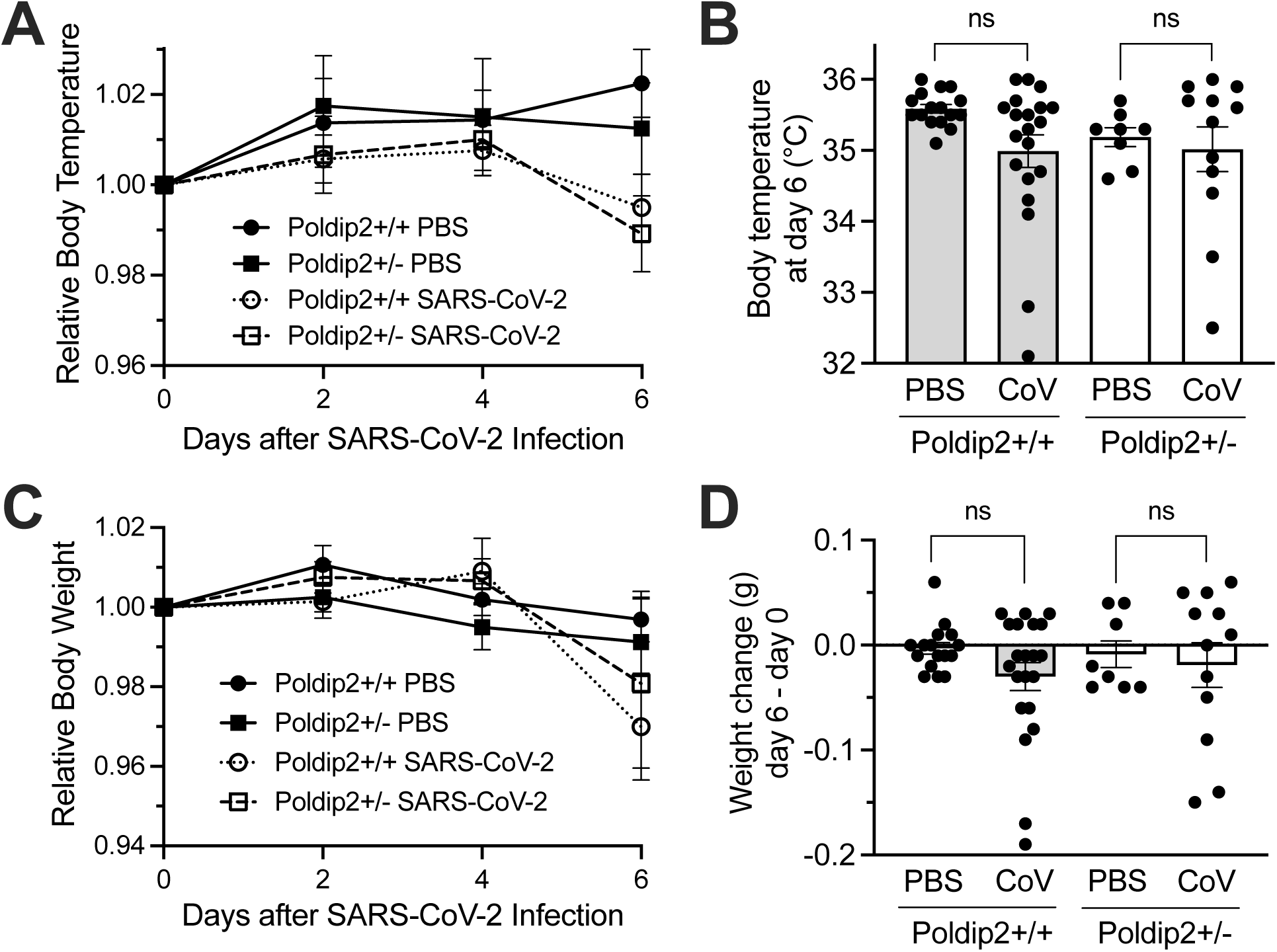
Acute infection does not significantly reduce body weights and temperatures. Body temperatures (A, B) and weights (C, D) were measured every other day after a single intranasal administration of PBS or SARS-CoV-2 to hACE2/Poldip2 mice. The graphs represent means ± SEM of data from n=8-21 animals. Time courses (A, C) were normalized to the initial value (day 0) for each mouse. Body temperatures at day 6 (B) and weight changes at day 6 relative to day 0 (D), were analyzed by 2-way ANOVA: ns, not significant.

### Poldip2 depletion reduces SARS-CoV-2-induced lung tissue damage

Given the previously described beneficial effect of reducing Poldip2 levels in a lipopolysaccharide (LPS)-induced acute lung injury model [18, 19], we sought to determine if Poldip2^+/-^ mice would exhibit similar effects in response to SARS-CoV-2 infection. Lung sections were collected at day 7 post-infection and processed for hematoxylin and eosin (H&E) staining. Histological evaluation revealed reduced lung tissue damage in Poldip2^+/-^ compared to Poldip2^+/+^ mice (**Figure 4**).

**Figure 4:**
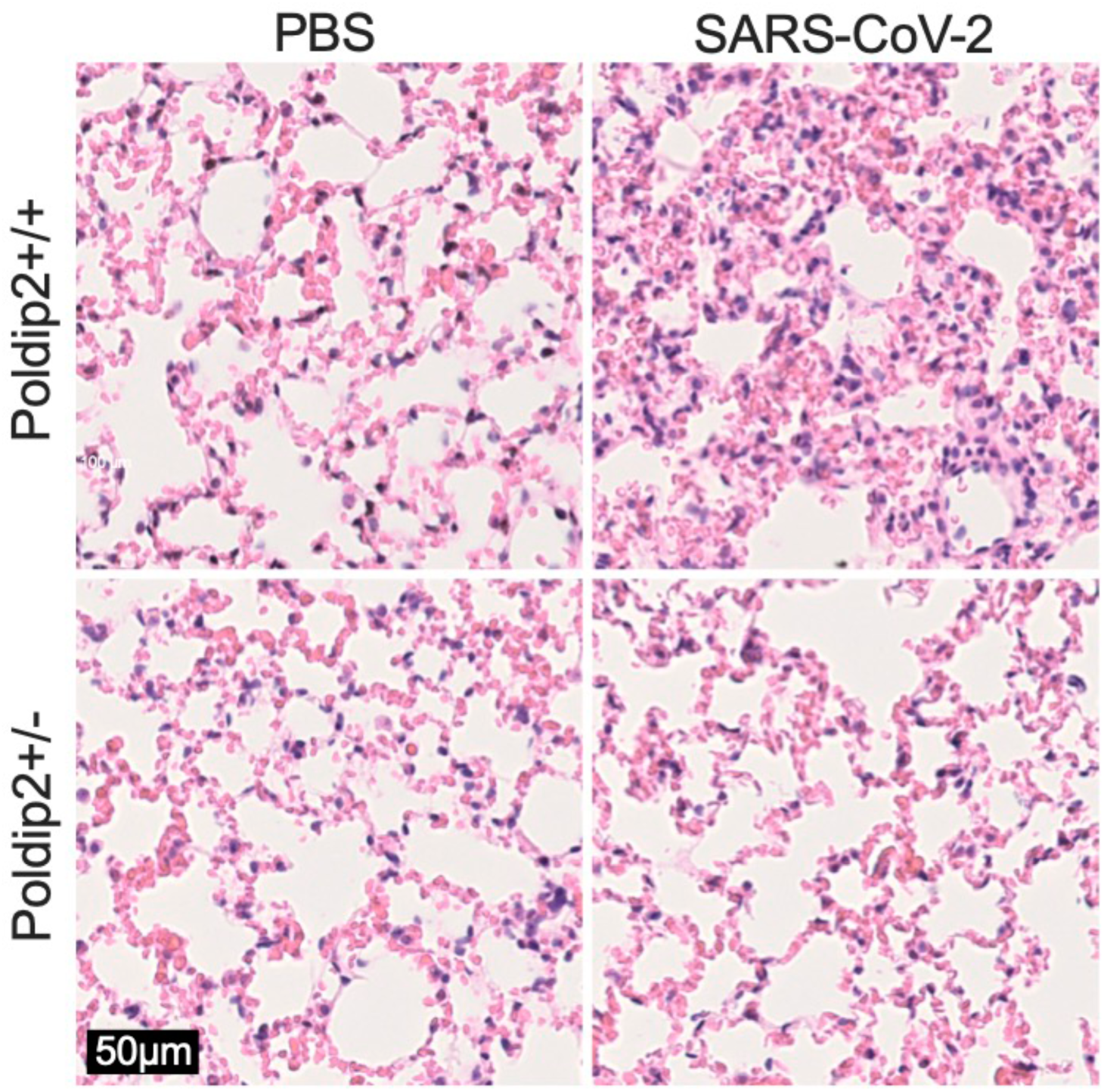
Poldip2 depletion alleviates lung tissue alterations induced by SARS-CoV-2 infection. hACE2/Poldip2 mice were sacrificed 7 days after intranasal administration of PBS or SARS-CoV-2. Lungs were processed for histology and paraffin sections were stained with hematoxylin/eosin. Images are representative of n=5-7 mice per group. Tissue damage and alveolar wall thickening resulting from SARS-CoV-2 infection in Poldip2^+/+^ mice (top right), were reduced in Poldip2^+/-^ animals (bottom right).

### Poldip2 mediates neutrophil infiltration in the lungs following SARS-CoV-2 infection

Since lung injury was reduced in Poldip2^+/-^ mice, we sought to investigate if Poldip2 is involved in neutrophil recruitment into the lungs after SARS-CoV-2 infection. Lung sections were collected at day 7 post-infection and processed for neutrophil-specific staining. Immunofluorescence data revealed a reduction in infiltrating CD45 and myeloperoxidase-positive neutrophils (**Figure 5**) in the lungs of Poldip2^+/-^ mice after SARS-CoV-2 infection, compared to Poldip2^+/+^ mice. This effect is consistent with the decrease in lung injury observed above in Poldip2 knockdown mice.

**Figure 5:**
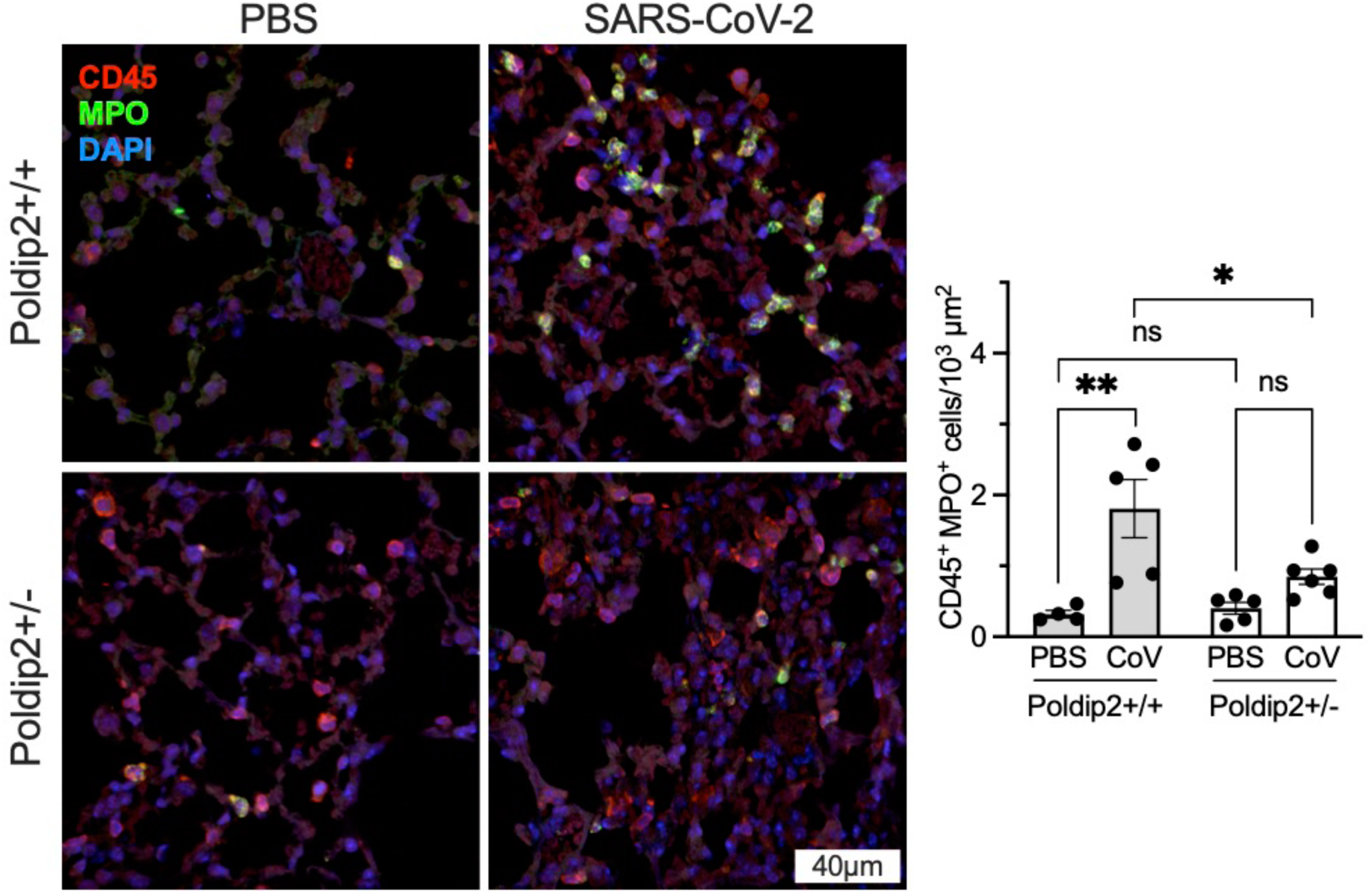
Poldip2 depletion reduces neutrophil infiltration in lung after SARS-CoV-2 infection. Lungs from hACE2/Poldip2 mice were harvested 7 days after infection and processed for immunofluorescence. Tissue sections were stained to detect the leukocyte common antigen (CD45, red), the neutrophil marker myeloperoxidase (MPO, green) and nuclei (blue), as shown in representative micrographs (left). Cells positive for both CD45 and MPO were counted in 10-13 images per animal (right). Bars represent means ± SEM of data from n=4-6 mice. Data were analyzed using 2-way ANOVA: ns, not significant; *P<0.05, **P<0.01.

### Poldip2 depletion reduces chemokines induced by SARS-CoV-2 lung infection

SARS-CoV-2 infection has been shown to be associated with increased expression of several chemokines. To determine whether Poldip2 regulates chemokine induction following SARS-CoV-2 infection, we first examined CXCL1 and MCP1 mRNA levels in lung tissue. Seven days after intranasal administration of SARS-CoV-2, both CXCL1 and MCP1 mRNA were significantly increased in the lungs of Poldip2^+/+^ mice. While no difference was observed in MCP1 levels between Poldip2^+/+^ and Poldip2^+/-^ mice, CXCL1 mRNA levels were reduced in Poldip2^+/-^ animals (**Figure 6A**). In contrast, Poldip2 depletion completely prevented increases in MCP1 protein levels in BAL, while CXCL1 protein expression was not induced in either Poldip2^+/+^ or Poldip2^+/-^ mice (**Figure 6B**). These data support the idea that upregulation of chemokines by Poldip2 may contribute to its stimulation of leukocyte recruitment after the viral infection.

**Figure 6:**
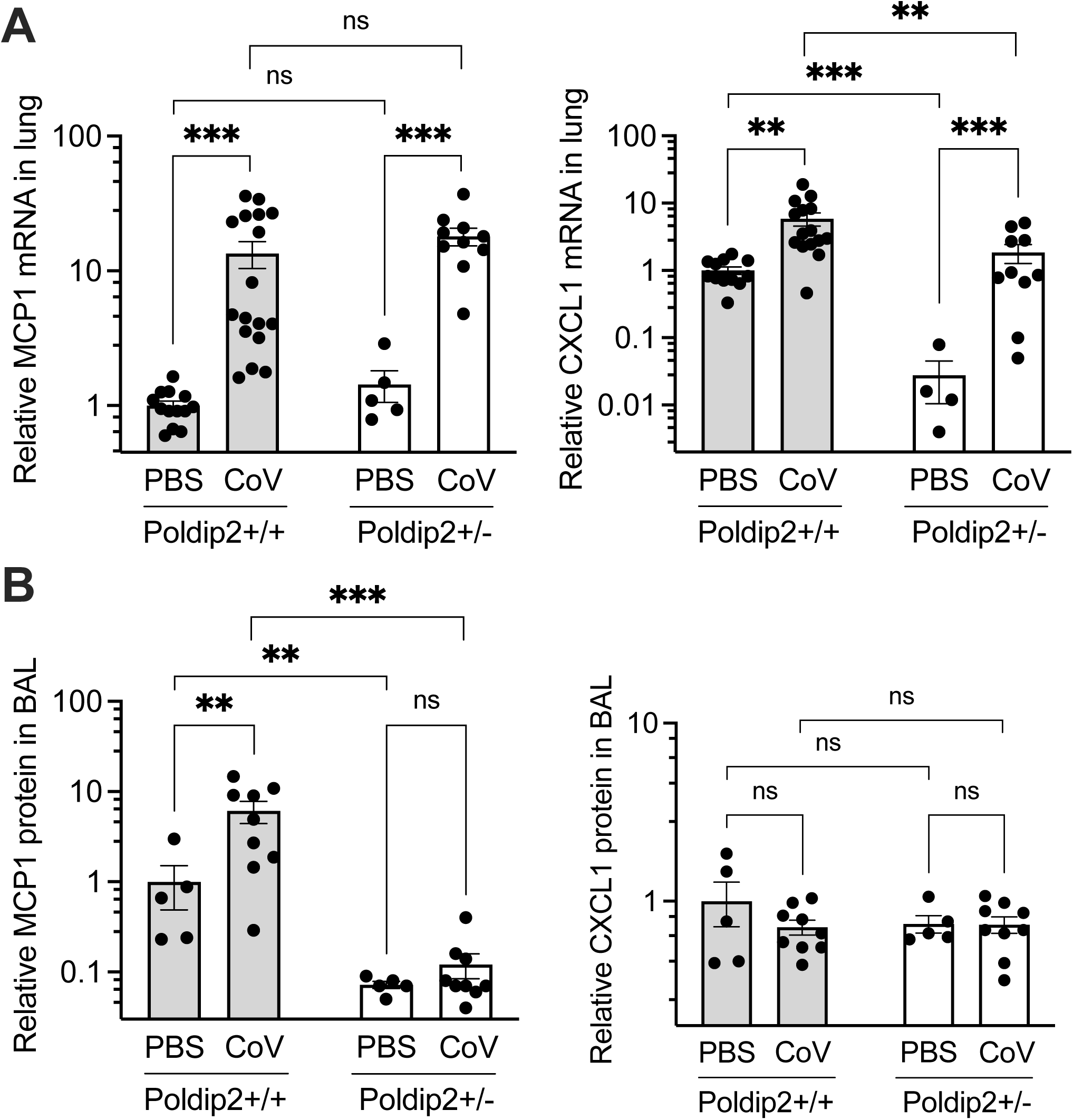
Poldip2 depletion reduces chemokine expression. hACE2/Poldip2 mice were sacrificed 7 days after intranasal administration of PBS or SARS-CoV-2. Bronchoalveloar lavage (BAL) fluid was collected before harvesting lungs for RNA extraction as in Figure 2. mRNAs were measured by RT-qPCR in lung tissue (A) and protein assays were carried out by ELISA in bronchoalveolar lavage fluid (B). Bars represent means ± SEM of data from n=4-17 animals. Data were analyzed using 2-way ANOVA: ns, not significant; ** P<0.01; *** P<0.001.

### Effects of Poldip2 depletion on cytokine induction by SARS-CoV-2 in lung and BAL fluid

To determine whether Poldip2 also regulates cytokine induction after SARS-CoV-2 infection, we first examined expression in lung tissue. TNFα, IL-1β, IL-6 and IFN-γ mRNA levels were measured 7 days after SARS-CoV-2 infection. TNFα and IFN-γ were significantly upregulated in Poldip2^+/+^ mice. However, their mRNA expression was not affected by Poldip2 depletion. Interestingly, upregulation of both IL-1β and IL-6 mRNAs were unexpectedly exacerbated in Poldip2^+/-^, compared to Poldip2^+/+^ mice in response to SARS-CoV-2 (**Figure 7A**). Additionally, TNFα, IL-1β, IL-6 and IFN-γ protein levels were evaluated in BAL fluids of Poldip2^+/+^ and Poldip2^+/-^ mice 7 days after SARS-CoV-2 infection using ELISAs. Corroborating the mRNA expression data, IFN-γ protein levels were increased in both Poldip2^+/+^ and Poldip2^+/-^ mice and no difference was found between the two genotypes. Similar results were observed for IL-6 protein levels. In contrast, TNFα, and IL-1β protein levels were increased 7 days after SARS-CoV-2 in BAL of Poldip2^+/+^ mice, but not in Poldip2^+/-^ animals (**Figure 7B**).

**Figure 7:**
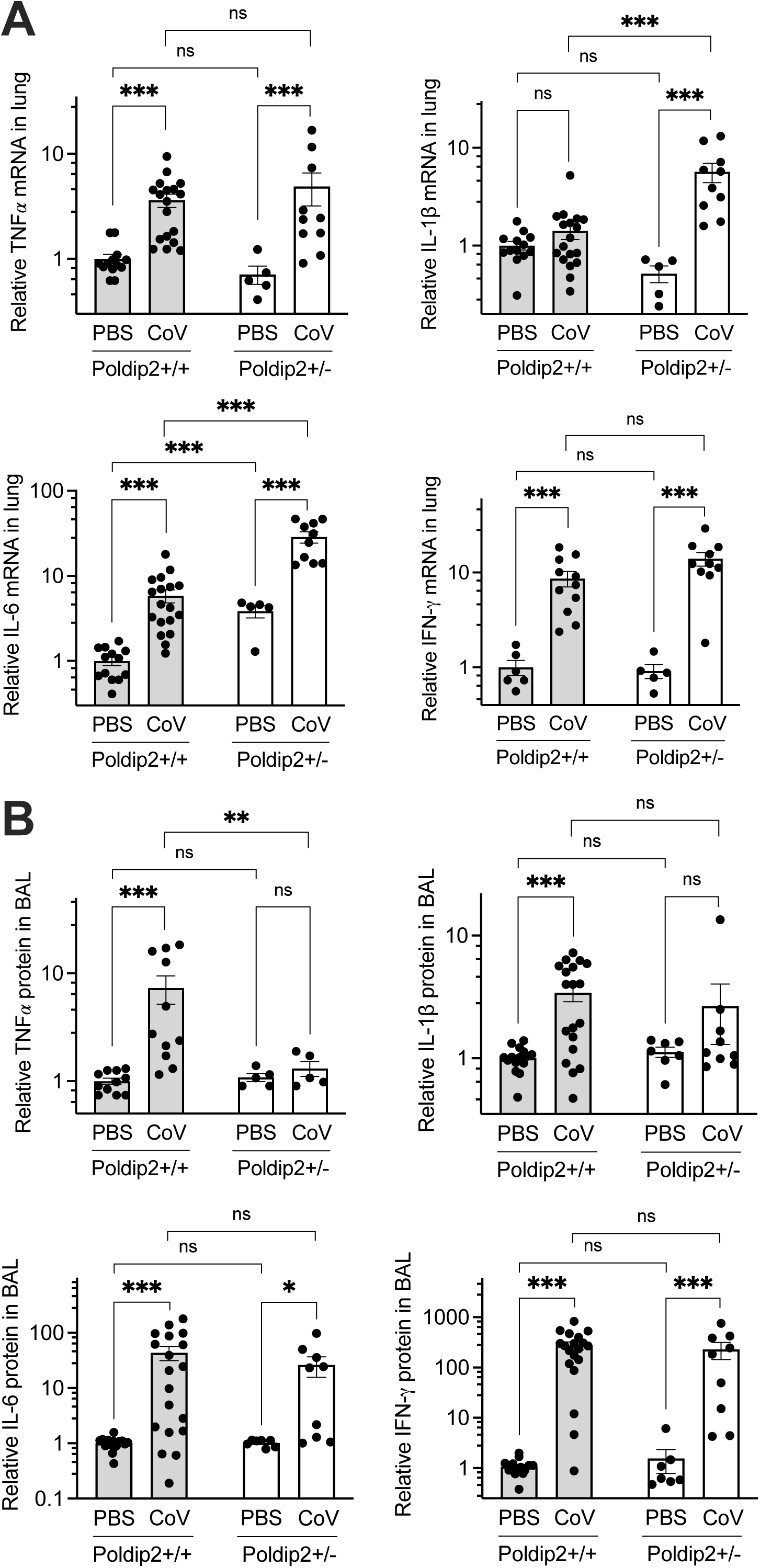
Poldip2 depletion selectively modifies cytokine expression after SARS-CoV-2 infection. Bronchoalveolar lavage fluid and lungs from hACE2/Poldip2 mice were collected 7 days after infection and processed as in Figure 6 to measure the expression of cytokine mRNAs (A) and proteins (B). Bars represent means ± SEM of data from n=5-19 animals. Data were analyzed using 2-way ANOVA: ns, not significant; * P<0.05; ** P<0.01; *** P<0.001.

## Discussion

The present study revealed that Poldip2 is upregulated in the lungs of COVID-19 patients and assessed the therapeutic potential of targeting Poldip2 in a mouse model. Poldip2 knockdown alleviated SARS-CoV-2-induced tissue damage, neutrophil infiltration and inflammation. The beneficial effects of Poldip2 depletion were most notable in BAL fluid, with blunted induction of MCP1, TNF𝛼 and IL-1β, following viral infection. Therefore, our data implicate Poldip2 in the inflammatory response triggered by SARS-CoV-2 and point to a potential novel therapeutic avenue.

In humans, SARS-CoV-2 infection presents a broad clinical spectrum ranging from asymptomatic to severe illness [28]. Several animal models that recapitulate the clinical and pathological characteristics of COVID-19 have become invaluable tools for elucidating the biological pathways underlying SARS-CoV-2 infection [2]. Because the spike protein of the initial SARS-CoV-2 strain fails to engage murine ACE2, in this study we used a transgenic mouse model in which the human ACE2 is overexpressed under the K18 epithelial promotor. Thus, human ACE2 is expressed in airway epithelial cells, colon and to a lesser extent kidney, liver, spleen, and small intestine [29, 30]. Therefore, these transgenic mice are susceptible to SARS-CoV-2 infection and develop acute respiratory disease after intranasal exposure, reproducing the major elements of severe disease observed in humans [30]. Previous studies using K18-hACE2 mice have demonstrated that intranasal exposure to 2 different doses of SARS-CoV-2, 2×10^3^ or 2×10^4^ PFU, resulted in acute disease with considerable weight loss (12%-20%) and lung injury, with some animals at the lower dose surviving infection despite significant weight loss [30]. A different study using the same animal model has shown that mice infected with 2.5×10^4^ PFU presented a 25% weight loss by day 7 after infection, which was less pronounced and delayed in mice infected with a lower dose, 1×10^2^ PFU [31], suggesting dose-dependent SARS-CoV-2 manifestations. In our study (using 4x10^5^ PFU), starting on day 4 Poldip2^+/+^ K18-hACE2 animals began to show signs of SARS-CoV-2 infection, presenting trends towards decreases in body temperature and weight loss, but to a lesser extent than in published studies. This difference may be due to reduced live virus after storage or to greater supportive care of experimental animals in our study. Likewise, in our hands, Poldip2^+/-^ mice exhibited a similar body temperature and weight loss tendency in response to SARS-CoV-2.

Our previous studies indicated an important role of Poldip2 in modulating cytokine and chemokine induction in an LPS-induced acute lung injury model [18, 19]. Heterozygous deletion of Poldip2 was protective against ARDS-induced TNFα, MCP1, IL-1β and CXCL1 in lung tissue. In addition, we recently found that endothelial-specific Poldip2 deletion *in vivo* also results in reduced expression of proinflammatory cytokines and chemokines including TNFα, CXCL1 CXCL2, IL-1β and IL-6 following LPS-induced ARDS [18]. Because Poldip2 seems to have such a profound effect in modulating the inflammatory response we hypothesized that Poldip2 depletion would also be protective after SARS-CoV-2 infection. In this study, SARS-CoV-2 led to a pronounced inflammatory response in lung tissue with upregulation of several pro-inflammatory cytokines and chemokines, including TNFα, IL-6, IL-1β, IFN-γ, MCP1, and CXCL1. Consistent with our previous studies using the LPS-induced ARDS model [19], we found that heterozygous deletion of Poldip2 significantly reduced TNFα, IL-1β, MCP1, and CXCL1 induction in BAL or lung tissue following SARS-CoV-2, indicating that Poldip2 knockdown led to an overall reduced inflammatory response. Interestingly, both IL-6, IL-1β mRNA levels were significantly upregulated in lung tissue of Poldip2^+/-^ mice when compared to Poldip2^+/+^ mice, suggesting that loss of Poldip2 has a complex modulatory effect on the inflammatory response.

Given the previously described effect of Poldip2 on leukocyte attachment and infiltration into inflamed tissues [17–19], we sought to investigate the contribution of Poldip2 in leukocyte recruitment into the lungs after SARS-CoV-2 infection. Several studies have reported massive immune cell infiltration in the lungs during severe COVID-19 infection, with enhanced recruitment of myeloid cells [31, 32] and changes in lymphoid cells, such as T cells [33] and natural killer cells [34]. However, we still lack comprehensive insights into the immunopathology of post–severe COVID-19 infection in lung tissue. Our data show that SARS-CoV-2 led to pronounced MPO-positive neutrophil infiltration into the lungs, which was largely prevented by heterozygous deletion of Poldip2. The lack of neutrophil recruitment upon Poldip2 depletion could be explained by the reduced basal levels of the chemoattractant CXCL1 which has been shown to induce neutrophil recruitment [35]. However, it is also possible that neutrophils from Poldip2^+/-^ mice are not fully attaching to the inflamed endothelium, as our previous study has shown that Poldip2 mediates β2-integrin activation during neutrophil recruitment to inflamed lungs [36]. Corroborating our data, previous studies have also demonstrated that infected K18-hACE2 mice presented a pronounced increase in MPO-positive neutrophils in lung tissue upon exposure to SARS-CoV-2 [30]. Neutrophil extracellular traps, which are indicative of neutrophil activation and can contribute to inflammation-associated lung damage, have also been described in lung autopsy samples [37], highlighting the importance of neutrophil recruitment as a potential therapeutic target in severe COVID-19.

Exacerbated immune responses play a major role in the pathophysiology of SARS-CoV- 2, leading to severe lung injury [38]. Altogether, our results show that Poldip2 depletion reduced the overall inflammatory response and decreased neutrophil infiltration following SARS-CoV-2 infection, which in turn resulted in attenuated SARS-CoV-2-induced lung tissue damage in Poldip2^+/-^ mice, as demonstrated by our histological evaluation. In summary, our study reveals a potential beneficial effect of Poldip2 depletion in SARS-CoV-2 infection and further adds to our understanding of the detrimental role of Poldip2 on endothelial barrier function under inflammatory conditions. In addition, the identification of molecular mechanisms driving severe pathogenic processes in the lung, may provide critical insight into the molecular and cellular features of COVID-19 pathogenesis.

## Acknowledgements

We are grateful to the Georgia Medical Examiner’s Office for providing human lung tissue samples. We kindly thank the Emory National Primate Research Center (ENPRC) Director Paul Johnson; Division of Animal Resources, especially Denyse Levesque, Jennifer McMillan, and Kathy Martinez-Kautz for providing support in animal care and proficiency guidance and assessment for ABSL-3 work. Additionally, we thank Kalpana Patel and Maureen Thompson for ABSL3 biosafety guidance, and Justin Harper and Mirko Paiardini for assistance with biosafety protocols.

---

**Supplemental Figure 1.**
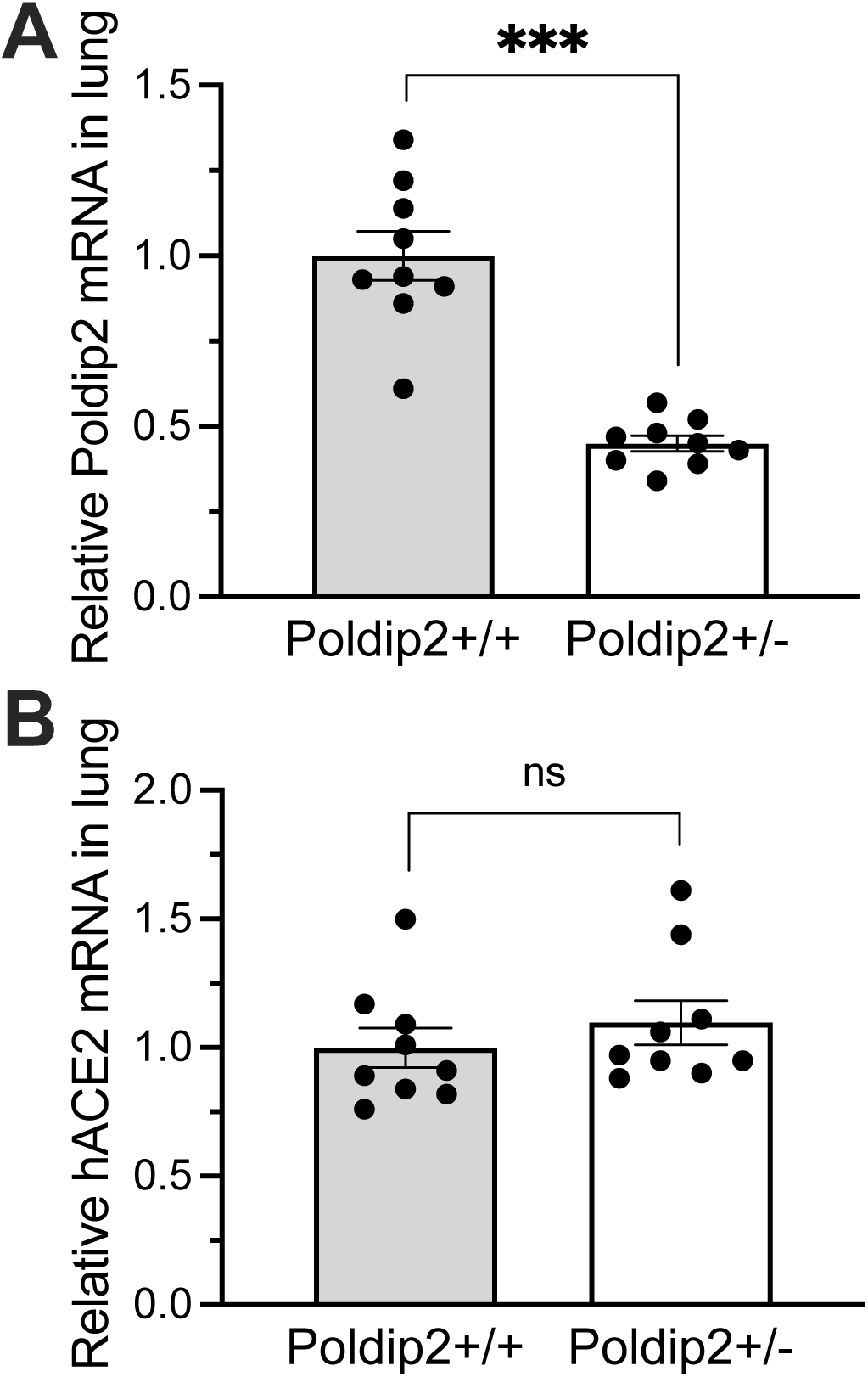
: Expression of mouse Poldip2 and human ACE2 mRNAs. Lungs were collected from hACE2/Poldip2 mice and processed to measure mRNAs by RT-qPCR as in Figures 6A and 7A. As expected, Poldip2 expression was reduced by about 50% in Poldip2^+/-^, compared to Poldip2^+/+^ controls (A). In addition, the expression of human ACE2 mRNA was not affected by the Poldip2 genotype (B). Bars represent means ± SEM of data from 9 animals. Means were compared using an unpaired two-tailed Student t-test: ns, not significant; *** P<0.001.

